# Evaluating batch correction methods for image-based cell profiling

**DOI:** 10.1101/2023.09.15.558001

**Authors:** John Arevalo, Ellen Su, Robert van Dijk, Anne E. Carpenter, Shantanu Singh

## Abstract

High-throughput image-based profiling platforms are powerful technologies capable of collecting data from billions of cells exposed to thousands of perturbations in a time- and cost-effective manner. Therefore, image-based profiling data has been increasingly used for diverse biological applications, such as predicting drug mechanism of action or gene function. However, batch effects pose severe limitations to community-wide efforts to integrate and interpret image-based profiling data collected across different laboratories and equipment. To address this problem, we benchmarked seven high-performing scRNA-seq batch correction techniques, representing diverse approaches, using a newly released Cell Painting dataset, the largest publicly accessible image-based dataset. We focused on five different scenarios with varying complexity, and we found that Harmony, a mixture-model based method, consistently outperformed the other tested methods. Our proposed framework, benchmark, and metrics can additionally be used to assess new batch correction methods in the future. Overall, this work paves the way for improvements that allow the community to make best use of public Cell Painting data for scientific discovery.

## Introduction

Image analysis has become a cornerstone of biological and biomedical research. Combining fluorescent labeling with advanced optical microscopy now allows us to visualize biological morphology, structures, and processes at unprecedented spatial and temporal resolution. Furthermore, high-throughput microscopy can now extract precise information about morphological changes caused by thousands of specific genetic or chemical perturbations. Analysis of the resulting image-based *profiles* – the typically thousands of measurements extracted from images of cells to capture their phenotype – can be used to deduce gene functions and disease mechanisms, as well as characterize mechanism and toxicity of potential therapeutics^1^. An image-based profile is a vector of values where each value corresponds to a particular morphological feature such as size, shape, intensity or texture of the cell. Image-based profiles are measured at the single-cell level but can be aggregated to the well or perturbation level, such that an experiment produces a very large matrix with rows as samples (cells, wells, or perturbations) and columns as features.

The most commonly used multiplex image-based profiling assay is Cell Painting^2,3^. Cell Painting uses six dyes to label eight cellular components (nucleus, nucleolus, endoplasmic reticulum (ER), Golgi, mitochondria, plasma membrane, cytoplasm, and cytoskeleton) that are imaged in five channels. Thus, each image-based profile captures rich morphological features that are extracted using automatic image processing and analysis pipelines. This approach offers single-cell resolution, captures valuable population heterogeneity, and provides distinct information from mRNA profiling ^4–9^ and protein profiling ^10^ at a low cost, with reagent costs of less than 25 cents per well and a yield of 1000-2000 single cells per well ^11^. Importantly, Cell Painting image-based profiles of cells exposed to different genetic or chemical perturbations have been successfully combined with machine learning strategies to generate predictive models that support key steps in drug discovery and development ^12^.

The broad applicability and the predictive power of Cell Painting data improves with the number of image-based profiles that can be used to either generate mechanistic hypotheses or build predictive models. Despite efforts from individual companies to create proprietary datasets, a large-scale, publicly available Cell Painting dataset is needed for the field to maximally advance. Other fields of biology, such as genomics, have proven the benefits of having a shared dataset in addition to shared goals.

Thus, to construct such a database, we recently partnered with colleagues from pharmaceutical companies, technology providers, and non-profit organizations to form the Joint Undertaking for Morphological Profiling (JUMP) Cell Painting Consortium^13^. These efforts resulted in the 2023 release of the first large-scale public dataset of image-based Cell Painting profiles, capturing data from more than 140,000 chemical and genetic perturbations (https://github.com/broadinstitute/cellpainting-gallery). A large, public dataset is most useful if it can be successfully queried using new profiles generated by individual laboratories in the future. With this in mind, the Consortium went to great lengths to embrace technical variation by exchanging compounds and generating images across twelve different laboratories which use varying pieces of equipment. This process created the opportunity to develop strong batch correction methods to align the data sources, which can then be used by future data generators.

The key challenge in aligning data across datasets is the presence of “batch effects” ^1^. In large-scale biological experiments, data is often collected in multiple batches, where a batch can refer to different experimental groups: multiple wells on a multi-well plate, multiple plates in a set processed in parallel, or multiple sets processed at a given laboratory. Batch effects refer to variations in the data that are not due to the biological variables being studied, but rather due to unintended technical differences across the experimental batches. These variations can arise from a multitude of factors such as different experimental conditions, processing times, or instrumentation used across separate batches of experiments. In the context of JUMP Cell Painting Consortium, technical factors (such as variations in the microscope used, microscope filters and settings, and cell growth conditions) all have effects on the image-based profiles we collected. Moreover, some batch effects occur even across batches from a single site, such as unintentional changes in lamp intensity, staining concentration, and cell seeding or growth rate. Additionally, alignment requires factoring in the hierarchical structure of Cell Painting experiments, with each readout originating from a region in a well within a multi-well plate, which in turn comes from an experimental batch at a particular laboratory.

Batch correction refers to methods which reduce batch effects, thus improving the ability to detect true biological signals. The definition of batch depends on the context of the data. In this paper, we consider two levels of batches: experimental batch, where multiple plates are produced simultaneously, and laboratory source, where multiple batches of data are produced by the same laboratory. Only a handful of batch correction methods have been developed and tested for image-based profiling. No systematic and comprehensive comparison and evaluation of such methods has been performed, making it unclear whether the available methods offer a reliable approach for dealing with batch effects in image-based profiling. Evaluations on single-cell RNA sequencing (scRNA-seq) batch correction methods reveal that no one method consistently outperforms the others ^14,15^ and suggest that most of these methods should not be used without the guidance of an expert ^16,17^. Therefore, it remains unknown whether any of the scRNA-seq batch correction methods can be reliably applied to image-based profiles.

Here, we carried out a comprehensive analysis of seven high-performing scRNA-seq batch correction methods, representing diverse approaches. We used qualitative visualizations along with four metrics that capture reduction in batch effects and six metrics that capture preservation of biological signals. We used the newly released public database created by the JUMP Cell Painting Consortium ^13^ to test the performance in the context of five common use cases: multiple batches from a single laboratory, multiple laboratories using the same microscope with few and many compounds, and multiple laboratories using different microscopes with few and many compounds. Given the practical constraints of working with large image-based profiling data, we focused our evaluation on population-averaged well-level profiles rather than single-cell level profiles. Population-averaged well-level profiles are computed by mean-averaging the morphological feature vectors for all cells in a well extracted with CellProfiler. We analyzed correction methods in the context of the replicate retrieval task (finding the replicate sample of a given compound across batches/laboratories), and we found that existing methods are effective in reducing batch effects in image-based profiles for some of the evaluated scenarios. Among the methods tested, Harmony, a non-linear method developed for processing scRNA-seq data, offered the best balance of removing batch effects and conserving biological variance. More broadly, the benchmark dataset, evaluation framework, and metrics we describe here will enable future assessment of novel batch correction methods as they emerge. Effective batch correction will advance the field and allow Cell Painting data to fulfill its potential for scientific discovery.

## Results

### Selection of batch correction methods and evaluation strategies

A major part of our work was to comprehensively survey methods for batch correction, as well as strategies for their evaluation. Given the rapid advancements in the field of scRNA-seq, particularly in the development of methods to address batch correction, we focused our attention on this area. We decided to test a subset of the better-performing methods identified in a recent analysis of scRNA-seq batch correction methods ^14,18^. These methods were available in Python and required no additional metadata. Additionally, the chosen methods were representative of different approaches and included linear methods (Combat ^19^ and Sphering ^20^), neural-network based methods (scVI ^21^ and DESC ^22^), neighbor-based methods (Scanorama ^23^ and MNN ^24^), and a mixture-model based method (Harmony ^25^).

We will briefly summarize the main characteristics of these methods, to enable the reader to place our results in the appropriate context. Combat ^19^ models batch effects as multiplicative and additive noise to the biological signal and uses a Bayesian framework to fit linear models that factor such noise out of the readouts. Sphering^20^ computes a whitening transformation matrix ^26^ based on negative controls and applies this transformation to the entire dataset. It requires every batch to include negative control samples for which variations are expected to be solely technical. scVI ^21^ is a variational autoencoder model for scRNA-seq data and learns a low-dimensional latent representation of each input that reduces the noise of batch effects. DESC ^22^ trains an autoencoder along with an iterative clustering algorithm to remove batch effects and preserve biological variation, and requires the knowledge of the biological variable of interest as input, which may be unknown at the batch correction stage. MNN ^24^ aligns representations between two batches by finding pairs of samples that are mutually nearest neighbors. Scanorama ^23^ computes mutual nearest neighbors across all of the batches and uses such neighbors as anchors to align the datasets. Harmony ^25^ is an iterative algorithm based on expectation-maximization that alternates between finding clusters with high diversity of batches, and computing mixture-based corrections within such clusters. All tested batch correction methods except Sphering require batch labels, Sphering alone requires negative control samples, and only DESC additionally requires biological labels. MNN, Scanorama, and Harmony necessitate recomputing batch correction across the entire dataset whenever new profiles are incorporated. While Sphering, Combat, scVI, and DESC don’t require recomputation, they don’t guarantee perfect corrections for new profiles from an unseen source.

Developed initially for scRNA-seq data, these methods also apply to morphological profiles, despite inherent biological and statistical differences. Most foundational assumptions about the methods, including the use of vector space metrics to reveal similarities, remain valid in the image-based profiling domain (See *Methods* section for assumptions of tested methods). However, we note a crucial difference in the manner we have applied these methods. Given the sheer volume of data in large image-based profiling datasets, which may contain billions of single cells compared to the millions typically found in scRNA-seq, it is computationally impractical to apply these methods at the single-cell level. Importantly three out of the seven methods require computing batch correction across the entire dataset. This means that subsampling, a strategy often used to manage large datasets, is not a viable option for those. Thus, we evaluated these methods based on their ability to correct batch effects in population-averaged well-level (or “pseudo-bulk”) profiles, rather than single-cell level profiles. Importantly, this shift does alter the distribution of the features, but we believe this is an acceptable trade-off given the computational constraints and the overall goal of correcting batch effects at a broader level.

Beyond batch correction methods for scRNA-seq, we also reviewed those specifically designed for image-based profiling data analysis. A small handful of past research in this domain incorporated a step for batch correction. Such approaches can be split broadly into two categories: those based on (a) pre-computed feature transformation and (b) representation learning.

Pre-computed feature transformation approaches learn a transformation of features that have been extracted from images. In such workflows, common normalization steps have been described by Bray and Carpenter ^27^ to deal with interplate and intraplate normalization - these aim to reduce local variances, but they are limited when technical variations are strong. Sphering ^20^ is the most used batch correction method for feature-transformation-based profiles widely applied in Cell Painting pipelines ^28,29^ and was included in our testing.

Next, we considered which dataset to use as a benchmark in our evaluation. RxRx1 ^30^ and RxRx3 ^31^ datasets are resources for image-based profiling, with millions of images associated with thousands of compounds, but derive from a single, highly-quality-controlled laboratory so cannot be used to assess methods aiming to correct more dramatic batch effects. We chose the JUMP Cell Painting Consortium data specifically because it originated from a diverse range of laboratories using different instruments and protocols, thus capturing the heterogeneity typical of large public datasets. Unlike other public resources that contain data from a single source, the diversity of data sources in the JUMP dataset provides a robust testbed to develop methods that can generalize to other heterogeneous datasets. It also allows mimicking a situation where an individual laboratory might attempt to align their data with public data collected in multiple laboratories.

We computed four metrics that report the effectiveness in removing batch effects and six metrics to measure how well the correction preserves biological information, previously reported in scRNA-seq benchmarks ^14,18^. The list of the 10 quantitative metrics we used in this study are described in the *Metrics* section. We also use UMAP ^32^ visualizations as a qualitative tool for assessing batch correction effectiveness.

### Scenario 1: Single microscope type, single laboratory, multiple batches, few compounds, multiple replicates

In this scenario, we analyzed 306 “landmark” compounds present on the *Target2* plates (see *Methods: Dataset description*), one of which is present in each of the 13 experimental runs/batches produced by a single laboratory (source_6). There were a median of 21 replicates per compound. Given that profiles were generated in the same laboratory, with many replicates and relatively low technical variance, this simplified scenario helped us establish a baseline for the best possible results while also guiding the pipeline that could be applied across all more complex scenarios. We considered the experimental run as the batch variable because it is the most dominant confounding source within a single laboratory’s data.

Image-based profiling requires carefully designing feature extraction and preprocessing pipelines to accurately represent the biological content of images. Preprocessing refers to the transformation and filtering steps that prepare raw data for in-depth analysis ^33^. Our preprocessing pipeline comprised four steps: (1) low-variance feature removal; (2) median absolute deviation normalization ^34^, which rescales plate-wise individual features; (3) rank-based inverse normal transformation ^35^, which transforms a variable into one that more closely resembles a standard normal distribution; and (4) feature selection, which removes redundant features using by correlation thresholding. See *Methods: Preprocessing pipeline* for details. We refer to this processed representation as the Baseline.

To quantify batch correction methods, we examined two kinds of metrics ^14,18^: (1) *batch removal* metrics that capture how well a method removes confounding variation from data; and (2) conservation of biological variance, termed here as *bio-metrics* for short, that capture the ability to preserve the variation related to a known biological variable (*e.g.* chemical compound in our case). There is a trade-off between bio-metrics and batch removal metrics. A method could remove all of the confounding batch variation but simultaneously destroy the biological signal in the data. Also notice that some of these metrics are sensitive to the number of samples per concept of interest (e.g. compound), others focus on a notion of local neighborhood, and others consider more global arrangement. Because different metrics capture different aspects of the correction procedure, and no individual metric captures both effects, we took them all into consideration when making comparisons between multiple batch correction methods. The average of such metric scores has been shown to be a reasonable ranking criterion ^14,18^. To make interpretation easier, every metric here was normalized between 0 and 1, with 0 being the worst performance and 1 being the best. These quantitative metrics (Sup Figure A) guided the selection of the preprocessing steps to be used as our Baseline approach, in agreement with established protocols in the field ^34,36^. All remaining results in this paper apply this four-step preprocessing.

Following the preprocessing, we applied the following batch correction methods that have previously been identified as top-performing methods when applied to scRNA-seq data: scVI ^21^, DESC ^22^, MNN ^24^, Scanorama ^23^, Combat ^19^, and Harmony ^25^. We reiterate that we used pseudo-bulk profiles and did not attempt batch correction at the single-cell level due to the computational time required to process up to billions of cells included in a typical image-based profiling experiment. Qualitatively, the 2D projection of the profiles after applying six different methods, in addition to our Baseline, shows no clusters associated with any particular batch (Sup Figure B). This suggests that all the methods were successfully mixing the profiles from different batches. However, we noticed that Harmony better grouped the data points associated with the same compound than others, suggesting that such methods are more effective at preserving biological information. Here, positive control compounds exhibited better clustering (see *Methods: Dataset description*), which was expected as these compounds were chosen based on the strong phenotype in previous Cell Painting experiments ^37^.

When evaluating the quantitative metrics (Table 1), all seven methods showed similar performance overall. In comparison to the Baseline, Combat and MNN achieved slightly better batch correction. Harmony surpassed the Baseline in bio-metric performance, particularly by increasing the mAP-nonrep score by 8%. This metric measures how effectively samples from the same compound can be retrieved from other non-replicate samples. Notably, scVI achieves the lowest overall batch correction score relative to the other methods.

**Table 1.** Evaluation Scenario 1. Quantitative comparison of seven batch correction methods measuring batch effect removal (four batch correction metrics) and conservation of biological variance (six bio-metrics). Metrics are mean aggregated by category. Overall score is the weighted sum of aggregated batch correction and bio-metrics with 0.4 and 0.6 weights respectively.

| Method | Batch correction |  |  |  | Bio metrics |  |  |  |  |  | Aggregate score |  |  |
| --- | --- | --- | --- | --- | --- | --- | --- | --- | --- | --- | --- | --- | --- |
|  | Graph connectivity | KBET | LISI batch | Silhouette batch | LISI label | Leiden ARI | Leiden NMI | Silhouette label | mAP (controls) | mAP (nonrep) | Batch correction | Bio metrics | Overall |
| Harmony | 0.79 | 0.42 | 0.40 | 0.74 | 0.98 | 0.07 | 0.60 | 0.55 | 0.95 | 0.60 | 0.59 | 0.62 | 0.61 |
| Combat | 0.78 | 0.44 | 0.41 | 0.84 | 0.98 | 0.05 | 0.56 | 0.53 | 0.94 | 0.55 | 0.62 | 0.60 | 0.61 |
| MNN | 0.78 | 0.44 | 0.40 | 0.84 | 0.98 | 0.07 | 0.56 | 0.52 | 0.94 | 0.54 | 0.61 | 0.60 | 0.61 |
| DESC | 0.78 | 0.39 | 0.42 | 0.74 | 0.98 | 0.06 | 0.58 | 0.55 | 0.95 | 0.56 | 0.58 | 0.61 | 0.60 |
| Baseline | 0.77 | 0.41 | 0.39 | 0.85 | 0.98 | 0.06 | 0.55 | 0.52 | 0.94 | 0.52 | 0.61 | 0.60 | 0.60 |
| Sphering | 0.74 | 0.39 | 0.37 | 0.87 | 0.98 | 0.06 | 0.54 | 0.52 | 0.94 | 0.51 | 0.59 | 0.59 | 0.59 |
| Scanorama | 0.54 | 0.72 | 0.53 | 0.59 | 0.98 | 0.09 | 0.56 | 0.45 | 0.94 | 0.49 | 0.59 | 0.59 | 0.59 |
| scVI | 0.72 | 0.39 | 0.43 | 0.70 | 0.98 | 0.06 | 0.58 | 0.53 | 0.94 | 0.56 | 0.56 | 0.61 | 0.59 |

In summary, Scenario 1 helped us optimize our evaluation pipeline, showing that the Baseline preprocessing produced good quality data for subsequent batch effect correction. Among the batch correction methods tested, Harmony, Combat, and MNN showed slightly better performance, but all methods were comparable.

### Scenario 2: Single microscope type, multiple laboratories, few compounds, multiple replicates

In this scenario we analyzed the *Target2* 306 compounds from 43 experimental runs/batches produced by three laboratories using the same model of microscope. We considered laboratory ID (identifier) as the batch variable because it is the most dominant confounding source (Sup Figure C). The Baseline approach was not able to integrate data from multiple laboratories, and batch effects were notable in the embedding, with clusters dominated by the confounding variable.

Unlike Scenario 1, Scenario 2 revealed variations in the efficacy of the methods when removing the confounding variable effect. Scanorama, Harmony, and scVI clustered samples from multiple laboratories more effectively than other methods. On the other hand, the Sphering correction did not improve the performance with respect to the Baseline. MNN, Combat and DESC did not differ significantly from the Baseline. When observing the embeddings labeled by compound (Sup Figure C), Scanorama, Harmony, and scVI were able to group samples from the same compound. On the other hand, the Baseline and Sphering showed clusters with Laboratory ID as the major separation criteria, creating one cluster per laboratory–compound pair.

Consistent with these qualitative observations, Scanorama and Harmony were also the top performer in the quantitative metrics (Table 2) for both batch removal and conservation of biological variance criteria. Taken together, we observed that by introducing stronger technical variations, i.e. analyzing data from multiple laboratories, the performance of batch effect correction methods decreased, compared to Scenario 1; however, methods’ rankings remained relatively consistent, with Harmony and Scanorama showing superior performance.

**Table 2.**
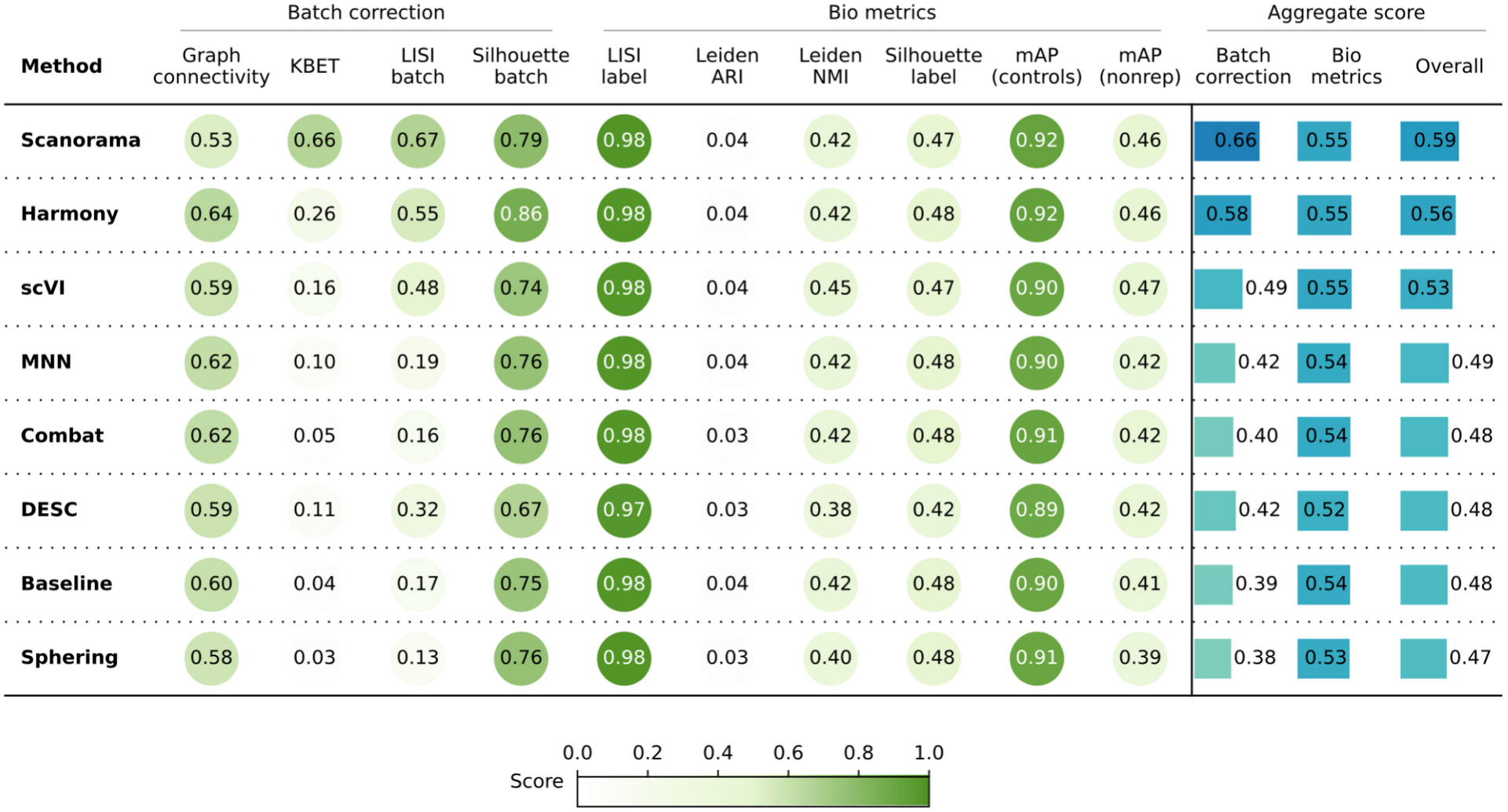
Evaluation Scenario 2. Quantitative comparison of seven batch correction methods measuring batch effect removal (four batch correction metrics) and conservation of biological variance (six bio-metrics). Metrics are mean aggregated by category. Overall score is the weighted sum of aggregated batch correction and bio-metrics with 0.4 and 0.6 weights respectively.

### Scenario 3: Single microscope type, multiple laboratories, multiple compounds, few replicates

In this scenario we analyzed 82,278 compounds from 43 experimental batches produced by three laboratories using the same model of microscope. We again used laboratory ID as the batch variable because it is the most dominant confounding source (Sup Figure D). This scenario posed an additional challenge due to the reduced number of replicates for most of the compounds; around 15,000 compounds had only one replicate, and ∼79,000 had 3 or fewer replicates. Importantly, the eight positive controls had around 2,500 replicates each.

Quantitatively (Table 3), Scanorama, Harmony, and scVI again obtained better results than the Baseline, Sphering, Combat, and MNN get comparable performances with the Baseline, and DESC underperformed in both bio-metrics and batch metrics. Overall, the increased complexity of the dataset resulted in a decreased gap across the methods. In other words, all methods struggled to remove the batch effects, as they remained notable after correction attempts. Compared to Scenario 2, the methods were generally less effective when dealing with more compounds and fewer replicates.

**Table 3.** Evaluation Scenario 3. Quantitative comparison of seven batch correction methods measuring batch effect removal (four batch correction metrics) and conservation of biological variance (six bio-metrics). Metrics are mean aggregated by category. Overall score is the weighted sum of aggregated batch correction and bio-metrics with 0.4 and 0.6 weights respectively.

| Method | Batch correction |  |  |  | Bio metrics |  |  |  |  |  | Aggregate score |  |  |
| --- | --- | --- | --- | --- | --- | --- | --- | --- | --- | --- | --- | --- | --- |
|  | Graph connectivity | KBET | LISI batch | Silhouette batch | LISI label | Leiden ARI | Leiden NMI | Silhouette label | mAP (controls) | mAP (nonrep) | Batch correction | Bio metrics | Overall |
| Scanorama | 0.54 | 0.33 | 0.36 | 0.76 | 1.00 | 0.01 | 0.29 | 0.27 | 0.20 | 0.05 | 0.50 | 0.30 | 0.38 |
| Harmony | 0.54 | 0.06 | 0.25 | 0.79 | 1.00 | 0.02 | 0.32 | 0.31 | 0.24 | 0.05 | 0.41 | 0.32 | 0.36 |
| scVI | 0.54 | 0.07 | 0.27 | 0.74 | 1.00 | 0.02 | 0.32 | 0.23 | 0.24 | 0.05 | 0.40 | 0.31 | 0.35 |
| Sphering | 0.54 | 0.02 | 0.00 | 0.80 | 1.00 | 0.00 | 0.26 | 0.35 | 0.21 | 0.03 | 0.34 | 0.31 | 0.32 |
| Combat | 0.54 | 0.03 | 0.02 | 0.76 | 1.00 | 0.01 | 0.29 | 0.30 | 0.22 | 0.04 | 0.34 | 0.31 | 0.32 |
| MNN | 0.54 | 0.03 | 0.01 | 0.75 | 1.00 | 0.01 | 0.29 | 0.29 | 0.20 | 0.04 | 0.33 | 0.30 | 0.32 |
| Baseline | 0.54 | 0.03 | 0.00 | 0.74 | 1.00 | 0.00 | 0.28 | 0.29 | 0.19 | 0.03 | 0.33 | 0.30 | 0.31 |
| DESC | 0.53 | 0.00 | 0.15 | 0.43 | 1.00 | 0.02 | 0.38 | 0.04 | 0.14 | 0.02 | 0.28 | 0.27 | 0.27 |

Score

### Scenario 4: Multiple microscope types, multiple laboratories, few compounds, multiple replicates

In this scenario we analyzed the *Target2* 306 compounds from 46 experimental runs/batches produced by five laboratories using three different high-throughput imaging systems. Three sources used the CellVoyager CV8000 system, one source used the ImageXpress Micro Confocal system, and one source used the Opera Phenix system. This Scenario was similar to Scenario 2 given the same number of *unique* compounds; however, in Scenario 4 the batch effects are mainly influenced by the differences in imaging technologies (Figure. 2.D) We again considered laboratory ID as the batch variable to stay consistent with Scenarios 2 and 3.

**Figure 1:**
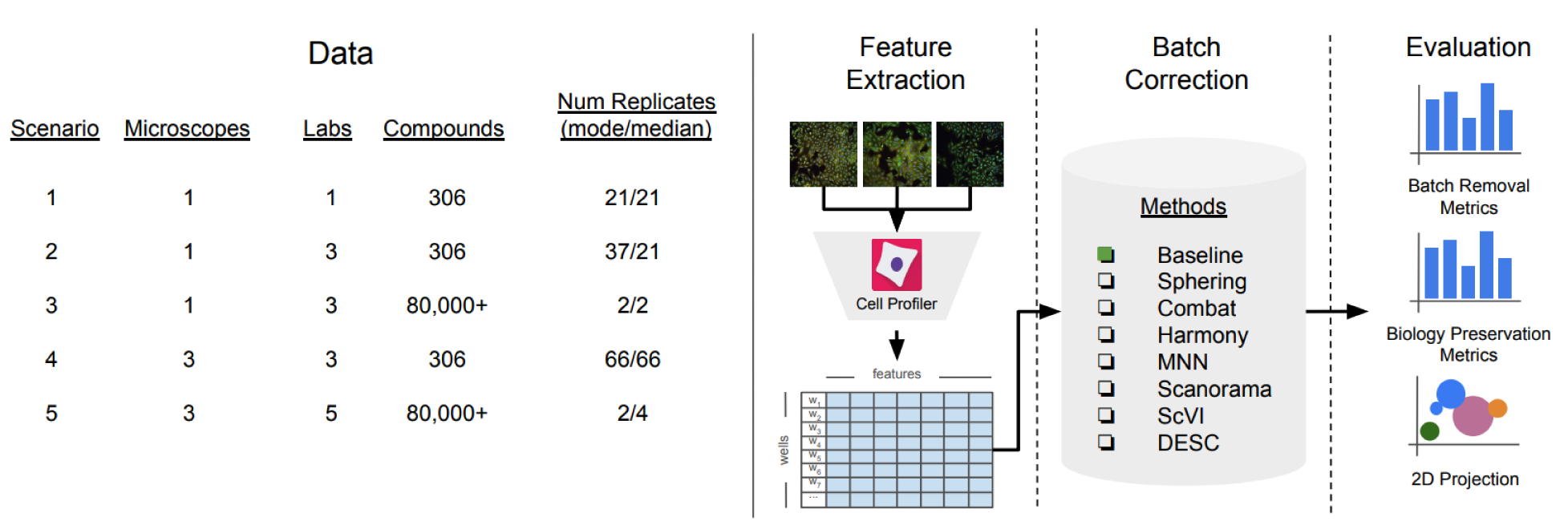
Evaluation pipeline. We evaluated five image-based profiling scenarios with different image acquisition equipment (high-throughput microscopes), laboratory, number of compounds and number of replicates. We used a state-of-the-art pipeline for image analysis. We compared seven batch correction methods using qualitative and quantitative metrics.

**Figure 2.**
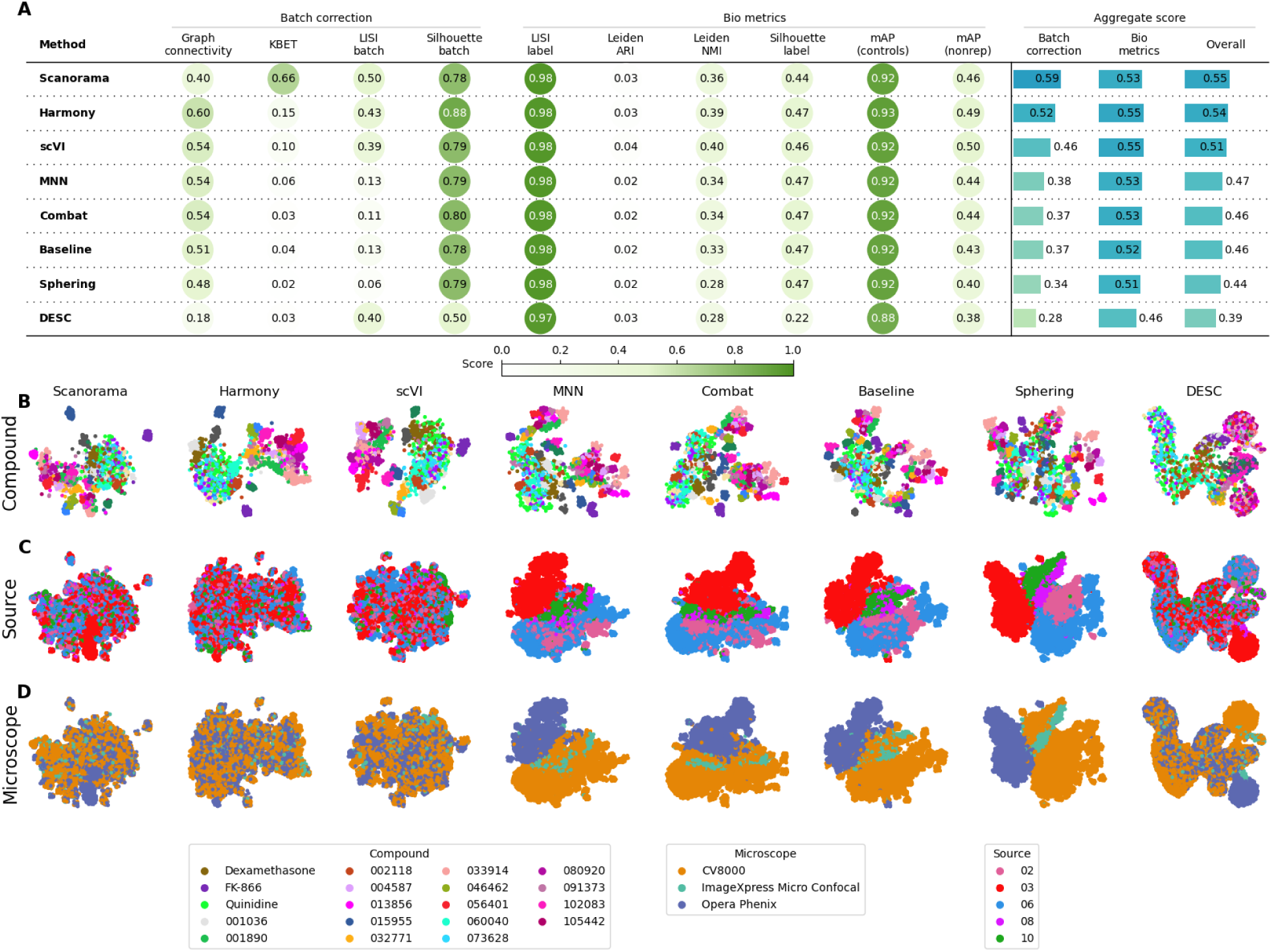
Evaluation Scenario 4. **A)** Quantitative comparison of seven batch correction methods measuring batch effect removal (four batch correction metrics) and conservation of biological variance (six bio-metrics). Metrics are mean aggregated by category. Overall score is the weighted sum of aggregated batch correction and bio-metrics with 0.4 and 0.6 weights respectively. Visualization of integrated data colored by **B)** Compound, **C)** Laboratory, and **D)** Microscope. Left-to-right layout reflects the methods’ descending order of performance. We selected 18 out of 306 compounds with replicates in different well positions to account for position effects that may cause profiles to look similar. Alphanumeric IDs denote positive controls.

We observed that Harmony and Scanorama were qualitatively better (Figure 2.B, 2.C), with Scanorama generating the best quantitative results (Figure 2.A). Similar to Scenario 2 and 3, MNN and Combat did not differ significantly from the Baseline. scVI outperformed linear models, and DESC and Sphering underperformed with respect to the Baseline. Compared to Scenario 2, the performance consistently decreased across methods and metrics, with the batch correction metrics exhibiting the highest drop. This indicates that the introduction of multiple microscope types had a strong impact in each method’s ability to align the data.

Compared to Scenario 2, the performance consistently decreased across all of the methods in most of the metrics, and the batch metrics exhibited the highest drop, indicating that differences in instrumentation had a strong impact on the methods’ ability to align the data.

### Scenario 5: Multiple microscope types, multiple laboratories, multiple compounds, few replicates

In the final, most complex scenario, we analyzed 82,412 compounds from 60 experimental runs/batches produced by five laboratories using different microscope systems as described in Scenario 4. We used laboratory ID as the batch variable. Again, we found that the differences between microscopes were the strongest confounding factor (Sup Figure E.C). The 2D embeddings for Combat, MNN and Sphering form several clusters; however those are still dominated by the source (Sup Figure E.B). Qualitative comparison helped us differentiate between methods that failed to integrate data from different sources and/or laboratories, but was less useful to check whether they preserved the biological information given the number of compounds being plotted in only two dimensions.

The quantitative results (Table 4) revealed that Scanorama outperformed other methods in batch correction metrics, whereas Harmony achieved the highest bio-metrics score. Compared to the Baseline batch correction scores, scVI showed marginal improvement, the linear methods performed similarly, and DESC underperformed. Notably, this scenario yielded the weakest overall performance, with a significant decline in batch correction scores compared to Scenario 3. This decline can be attributed to the integration of data from diverse microscope systems along with a large number of unique compounds.

**Table 4.**
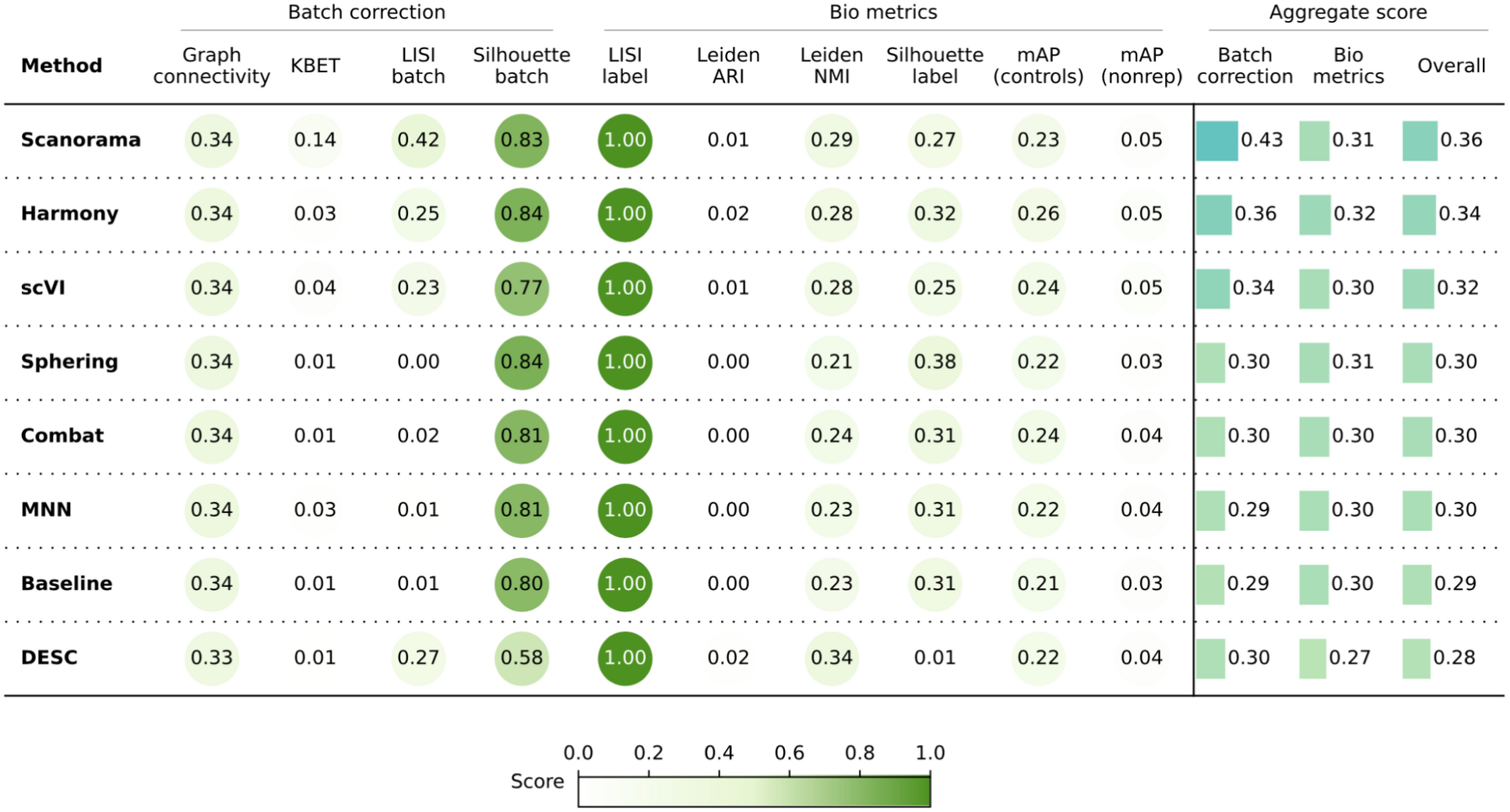
Evaluation Scenario 5. Quantitative comparison of seven batch correction methods measuring batch effect removal (four batch correction metrics) and conservation of biological variance (six bio-metrics). Metrics are mean aggregated by category. Overall score is the weighted sum of aggregated batch correction and bio-metrics with 0.4 and 0.6 weights respectively.

| Method | Batch correction |  |  |  | Bio metrics |  |  |  |  |  | Aggregate score |  |  |
| --- | --- | --- | --- | --- | --- | --- | --- | --- | --- | --- | --- | --- | --- |
|  | Graph connectivity | KBET | LISI batch | Silhouette batch | LISI label | Leiden ARI | Leiden NMI | Silhouette label | mAP (controls) | mAP (nonrep) | Batch correction | Bio metrics | Overall |
| Scanorama | 0.34 | 0.14 | 0.42 | 0.83 | 1.00 | 0.01 | 0.29 | 0.27 | 0.23 | 0.05 | 0.43 | 0.31 | 0.36 |
| Harmony | 0.34 | 0.03 | 0.25 | 0.84 | 1.00 | 0.02 | 0.28 | 0.32 | 0.26 | 0.05 | 0.36 | 0.32 | 0.34 |
| scVI | 0.34 | 0.04 | 0.23 | 0.77 | 1.00 | 0.01 | 0.28 | 0.25 | 0.24 | 0.05 | 0.34 | 0.30 | 0.32 |
| Sphering | 0.34 | 0.01 | 0.00 | 0.84 | 1.00 | 0.00 | 0.21 | 0.38 | 0.22 | 0.03 | 0.30 | 0.31 | 0.30 |
| Combat | 0.34 | 0.01 | 0.02 | 0.81 | 1.00 | 0.00 | 0.24 | 0.31 | 0.24 | 0.04 | 0.30 | 0.30 | 0.30 |
| MNN | 0.34 | 0.03 | 0.01 | 0.81 | 1.00 | 0.00 | 0.23 | 0.31 | 0.22 | 0.04 | 0.29 | 0.30 | 0.30 |
| Baseline | 0.34 | 0.01 | 0.01 | 0.80 | 1.00 | 0.00 | 0.23 | 0.31 | 0.21 | 0.03 | 0.29 | 0.30 | 0.29 |
| DESC | 0.33 | 0.01 | 0.27 | 0.58 | 1.00 | 0.02 | 0.34 | 0.01 | 0.22 | 0.04 | 0.30 | 0.27 | 0.28 |

Score

## Discussion

High-throughput image-based assays represent powerful strategies for making biological discoveries and facilitating development of new therapeutics. These assays, such as Cell Painting, capture large amounts of data that can be used to connect genetic or pharmacological perturbations to specific changes in cellular morphology and/or phenotype. Over the last decade, the amount of image-based data has grown exponentially. However, the data benchmarking, processing, management, comparison and evaluation of image-based profiles remains challenging. One of the current challenges for the field has been the lack of robust batch correction methods, making it difficult to compare image-based profile data from different instruments, laboratories or even between different batches from the same laboratory. Without such solutions, public databases will be useless. Here, we address this problem by creating a framework for evaluation and systematic comparison of batch correction methods for image-based profiling. We applied our strategy to comparing seven different batch correction methods that have been originally developed for use with scRNA-seq data.

Overall, across five relevant scenarios of increasing complexity that we tested (batches within a lab, across laboratories and across imaging instrumentation, all with more or fewer compounds), we found that Harmony and Scanorama consistently outperforms the other tested methods. We consider Harmony to be a good trade-off obtaining top-2 best performance in batch correction metrics, and the best performance in bio-metrics for all the scenarios we tested. In less complex scenarios, simpler methods may suffice; for example, for data generated in the same laboratory with many replicates of the compounds, the Baseline was sufficient to correct most of the batch effects, even though Harmony and Combat performed slightly better. However, we observed that as technical and biological variance increases, linear models such as Combat or Sphering fall short in aligning the data.

Although Harmony and Scanorama outperformed other methods tested, it’s important to point out a possible limitation for extensibility: it requires processing of all the data to re-align new batches against an existing dataset. This has a major impact on the ability of users to align their data with a large public dataset as it requires reprocessing and modifying existing representations. Therefore, further strategies, such as domain adaptation techniques ^38,39^ and self-supervised methods tailored for tabular data ^40^ may represent promising alternatives. These methods, grounded in machine learning principles, are inherently flexible, allowing for incremental learning and the integration of new data without altering the previously established representations. While still in the exploratory stages for our context, these approaches have demonstrated potential in similar high-dimensionality scenarios and may provide an effective solution to batch alignment challenges that arise when incrementally expanding large datasets ^41,42^.

Importantly, we discovered that in the most difficult-to-align scenarios, when there are more than a few hundred compounds and more than one microscope type, none of the methods are able to adequately remove the batch effects. In fact, the best methods provide the greatest improvement in the least difficult-to-align scenarios. This raises a call for advancements from the field. Our study focused on bulk (population-averaged) profiles. It is possible that applying batch correction at the level of single-cell profiles (likely subsampled), or even at a lower level using raw pixels, may yield better results. Among the methods we tested, Combat, DESC, and scVI are suited for training on subsampled data at the single-cell level. Although neural networks have been explored, implementation and evaluation of these methods at the JUMP-dataset scale with billions of cells still represents a challenge. Methods evaluated in this benchmark process tabular data extracted with standard image processing algorithms ^43^. An alternative approach is to learn representations from images ^29,36^. The design of neural network architectures and representation learning algorithms for Cell Painting is still an active research area, as is studying the interaction of these methods with those for batch correction; thus, our benchmark serves to establish a baseline for future studies comparing the performance of batch correction methods for learning-based representations. Finally, investigating quality control techniques, in addition to those employed in this study, may further enhance batch correction for methods that are sensitive to outliers. In downstream tasks, well-level representations are not considered in isolation; instead, we aggregate the five replicates of the same perturbation, with each replicate in a different batch, by computing the median profile. This way an outlier batch is less prone to contaminate results.

## Methods

### Dataset description

The JUMP Cell Painting dataset ^13^ is a collection of several datasets that were either generated or reprocessed by the JUMP Cell Painting Consortium. The primary dataset (*cpg0016-jump,* referred to as *cpg0016* for brevity) was generated during the data production phase of the JUMP-CP project. The compound dataset in *cpg0016* – referred to as *cpg0016[compound]* was generated across 13 data producing sites (or “sources”) and comprises a compound dataset (116,753 perturbations), an Open Reading Frame gene overexpression dataset (15,142 perturbation) and a CRISPR gene knockout dataset (7,977 perturbations).

A positive control plate of 306 diverse compounds – named *JUMP-Target-2-Compound* – was run with every batch of data generation. These plates not only allow alignment of data within the JUMP dataset, but also with future datasets generated outside the consortium and can thus be considered as control plates. In this paper, *JUMP-Target-2-Compound* plates are referred to as *Target2* plates for brevity; the remaining plates – comprising the 116,753 chemical perturbations are referred to as *Production* plates.

All Production plates have negative controls and several positive controls to identify and/or correct for different experimental artifacts. Dimethyl Sulfoxide (DMSO) -treated wells serve as *negative controls*. They are used for detecting and correcting plate to plate variations. They can also be used as a baseline for identifying perturbations with a detectable morphological signal.

### Metrics

We surveyed various evaluation strategies and considered them for the context of image-based profiling batch correction. In contrast to the scRNA-seq domain where multiple benchmark studies exist ^14,18^, image-based profiling lacks a comprehensive comparison of batch effect correction methods, and there are no agreed evaluation criteria.

Quantitative evaluation of batch correction methods broadly take two forms: using bespoke metrics to directly quantify batch effect removal and conservation of biological variance, or indirectly examining the expressivity of corrected image-based profiles through subsequent analysis tasks. An indirect approach is to utilize downstream tasks such as image classification ^44^ and perturbation replicate retrieval across batches ^45^ as a measure of the expressivity of the image-based profiles after correction. Used in isolation, this indirect approach may fail to detect the presence of batch effects in situations where the experimental covariates (e.g. plate ID) correlate with the labels of the downstream task (e.g. samples’ replicates).

Sypetkowski et al. ^30^ proposes a direct approach with their batch generalization and batch classification accuracy metrics. Batch generalization utilizes a perturbation classifier to compute the performance ratio between batches used for training the classifier and a held-out set of batches. A high ratio indicates a corrected representation that’s not heavily influenced by the batch. Batch accuracy trains a classifier to predict the batch ID based on the corrected representation; batch accuracy is low if batch correction has performed well. However, this approach also requires the tuning of additional classification pipelines during evaluation, making the comparison of batch correction methods somewhat less straightforward. Lastly, Kim et al.’s ^36^ evaluation provides another direct approach, using silhouette score, graph connectivity, and local inverse Simpson’s index (LISI) to quantify batch effects across different representation learning methods. Their evaluation is similar to our Scenarios 1 and 2.

We computed four metrics for batch removal efficiency and six metrics to measure how well the correction preserves biological information, previously reported in scRNA-seq benchmarks ^14,18^. They are implemented in the scib package^1^. To make interpretation easier, every metric is normalized between 0 and 1, with 0 being the worst performance and 1 being the best. A short description of each method is provided below. Mathematical definitions are provided in Luecken et al. ^18^.

- *ASW-based*: The average silhouette width (ASW) is a measure of how well a sample is assigned to its cluster. **Silhouette Label** metric considers the compound as the cluster ID; while **Silhouette Batch** metric considers the confounding variable as the cluster ID.
- **Graph_conn**: Uses the k-NN graph to measure the connectivity of each sample and those that belong to the same compound. This metric assumes If the batch effects were effectively removed, then the elements of the same biological concept should be close together.
- *LISI-based*: LISI is a metric based on the Simpson’s diversity Index to measure the diversity of a sample’s local neighborhood in the data. **LISI batch** uses the confounding variable to measure such diversity; while **LISI label** uses the compound annotations to measure such diversity. Variability in LISI label scores is <1e-3 for most of the scenarios. We confirmed this high-value low-variance behavior is also present in the scRNA-seq benchmarks^18^. We keep it for the completeness of its counterpart LISI batch.
- **kBET:** Compares the global distribution and the local distribution of the confounding variable for each sample in the dataset. If the confounding variable is effectively removed, then such distributions should be similar.
- **Leiden ARI** *and* **Leiden NMI**: Adjusted Rand Index (ARI) and Normalized Mutual Information (NMI) measure the agreement between two clustering assignments. **Leiden ARI** and **Leiden NMI** metrics computes agreement between the clustering assignments of the Leiden algorithm applied over the corrected data and the compound annotations.

We additionally report **mean average precision (mAP)** as a bio-metric (how distinguishable are samples of the same compound from other compounds – i.e., are they retrieved towards the top of a list of samples ranked by similarity to the query compound?). We measure similarity between samples using cosine similarity.

Following the information retrieval convention, each sample in the dataset is considered a query; M-1 other samples sharing the same compound are considered positive elements to be retrieved; and N samples comprising either (1) wells from the same plate as the query but treated with a different compound (mAP nonrep), or (2) the negative controls from the same plate (mAP control), are considered negative elements. For each query, a rank list is computed using the cosine similarity between the query profiles and the remaining (M-1)+N profiles. This ranked list is scored using average precision ^46^, which assesses the probability that positive elements will rank highly on the list. *AP* of the *i^th^* query can be expressed via relative change in recall:

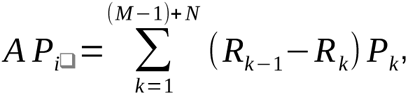 where

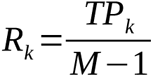 is recall at rank *k* (recall@k)

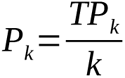 is precision at rank *k* (precision@k),

*TP_k_* is the number of all positive elements retrieved up to rank *k*.

Finally, we average the AP across all replicates of the compound, termed **mAP** for that compound.

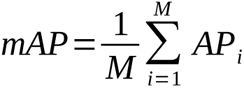

### UMAP visualization

We utilize UMAP visualizations as a qualitative tool for assessing batch correction effectiveness. While we acknowledge the issues associated with UMAP, as highlighted in recent studies showing how extreme dimensionality reduction can significantly distort high-dimensional data ^47^, we still employ UMAP visualizations for qualitative assessment of batch correction. This approach is useful for visual analysis: clusters corresponding to biological characteristics suggest successful correction, but clusters aligned with batch variables may indicate inadequate correction. Nevertheless, we recognize the need for caution in interpreting these visualizations due to the potential distortions inherent in such dimensionality reduction techniques.

### Distributional assumptions of tested batch correction methods

The batch correction methods tested in this work make different assumptions about the data distributions:

- MNN: Assumes batch effects are orthogonal to the biological information.
- scVI: Assumes negative binomial distribution for the features.
- Harmony: Assumes batch effects can be removed by iterative linear transformations. There are no assumptions on feature distributions.
- Combat: Assumes features are normally distributed and batch effects are multiplicative and additive noise^19^.
- Scanorama: As this method extends MNN by “stitching” together multiple datasets based on the mutual nearest neighbors, it similarly assumes batch effects are orthogonal to the biological information.
- DESC: Assumes technical differences across batches are smaller than true biological variations. There are no assumptions on feature distributions.
- Sphering: Assumes that negative controls sampled from different batches ought to be similar to each other in the biological sense, and any deviations from this normal-looking phenotype are rather technical. We applied sphering as described in Caicedo et al.^8^.

### Preprocessing pipeline

The 4762 cell profiler features measure shape, color, texture, and pixel statistics. Some of these measurements vary a lot in scale, distribution, while others are mostly constant across the dataset. Thus preprocessing is required. We explored combinations of processing steps (*Supplementary material: Preprocessing exploration*), included strategies to deal with outliers, and settled on the four steps listed below, applied in order.

**1) Variation filtering** The first step is to filter out features with low variance. We defined the absolute coefficient of variation *Cvar* of a feature *X* as:

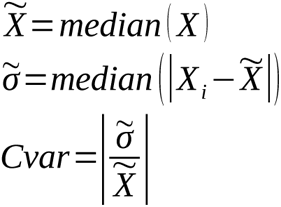 We compute *Cvar* for every feature plate-wise using control wells only. We discard features with any *Cvar* <1 *e*^−3^.
**2) Median absolute deviation** For every plate, we compute 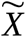 and 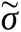 using the control wells only. Then for every well in a plate we transform the feature values as follows:

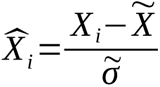
**3) Rank-based Inverse normal transformation (INT)**

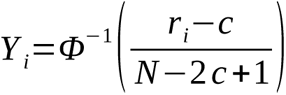
**4) Feature selection** We select features using Pycytominer ^48^. For any pair of features, if their correlation is higher than 0.9, then it will exclude the feature with the highest sum of correlation with the other features in the dataset.

## Acknowledgments

The authors gratefully acknowledge a grant from the Massachusetts Life Sciences Center Bits to Bytes Capital Call program for funding the data production. We appreciate funding to support data analysis and interpretation from members of the JUMP Cell Painting Consortium and from the National Institutes of Health (NIH MIRA R35 GM122547 to AEC). The authors appreciate helpful comments from Beth Cimini, Niranj Chandrasekaran, Arnaud Ogier, Thierry Dorval, and Jeremy Grignard.

## Author contributions

J.A. wrote the code and conducted the analysis; E.S. and R.V.D. performed exploratory analyses, S.S. and A.E.C. supervised the research. All authors designed the experiments and wrote and edited the paper.

## Data and code availability

All code to reproduce this analysis is provided as a reproducible Snakemake ^49^ pipeline at https://github.com/carpenter-singh-lab/2023_Arevalo_BatchCorrection. All the corresponding data is available as part of the *cpg0016-jump* dataset ^13^, available from the Cell Painting Gallery on the Registry of Open Data on AWS (https://registry.opendata.aws/cellpainting-gallery/).

## Supplementary material

### Preprocessing exploration

We explored combinations of the four steps above along with strategies to deal with outliers such as imputation (with KNN and median), clipping, and feature dropping. We choose the most convenient pipeline based on the mAP scores in Scenario 1. See https://github.com/carpenter-singh-lab/2023_Arevalo_BatchCorrection/issues/4 for further details about the exploration.

**Supplementary Figure A.**
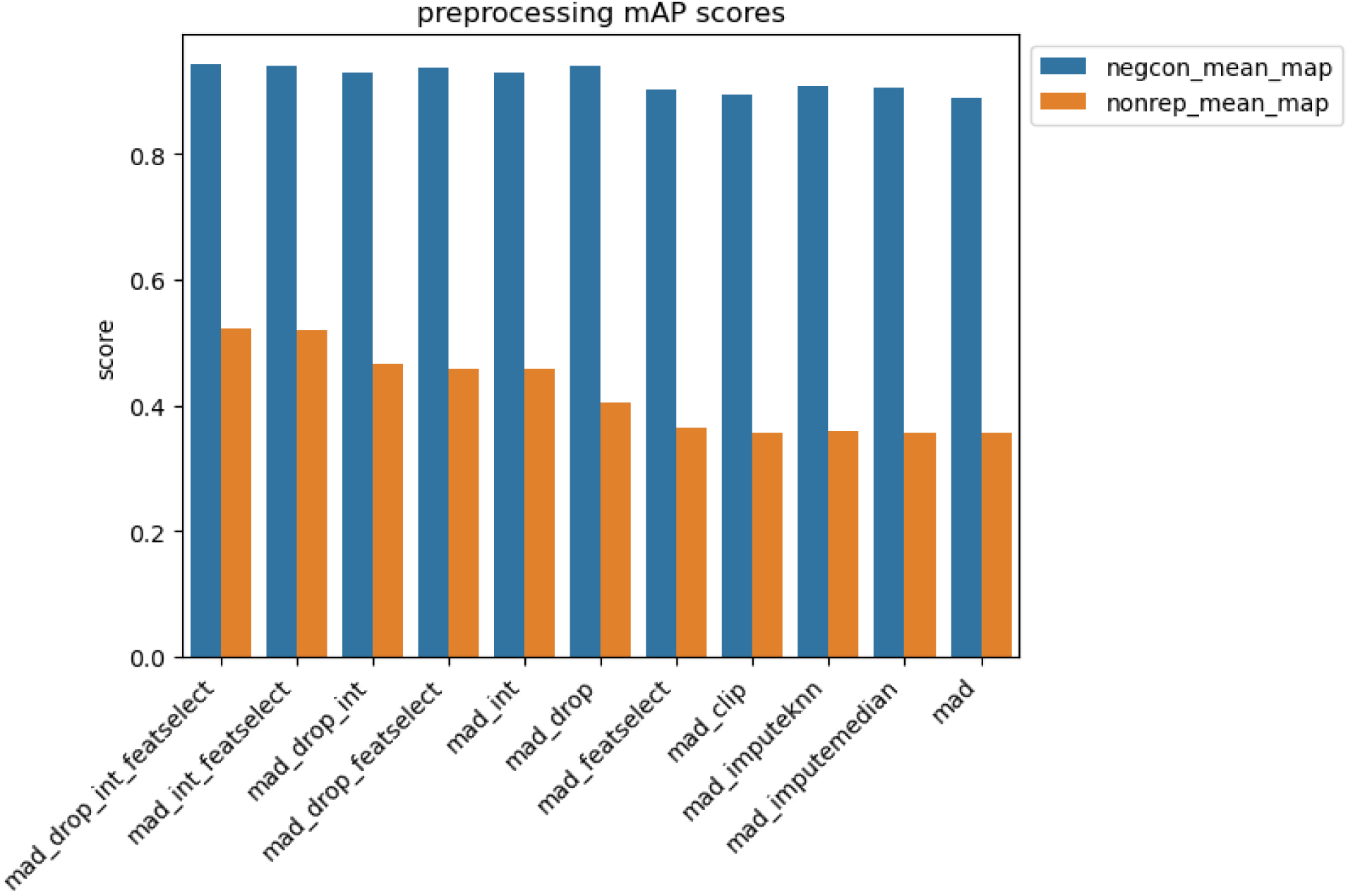
preprocessing mAP scores

### Scenario 1

**Supplementary Figure B.**
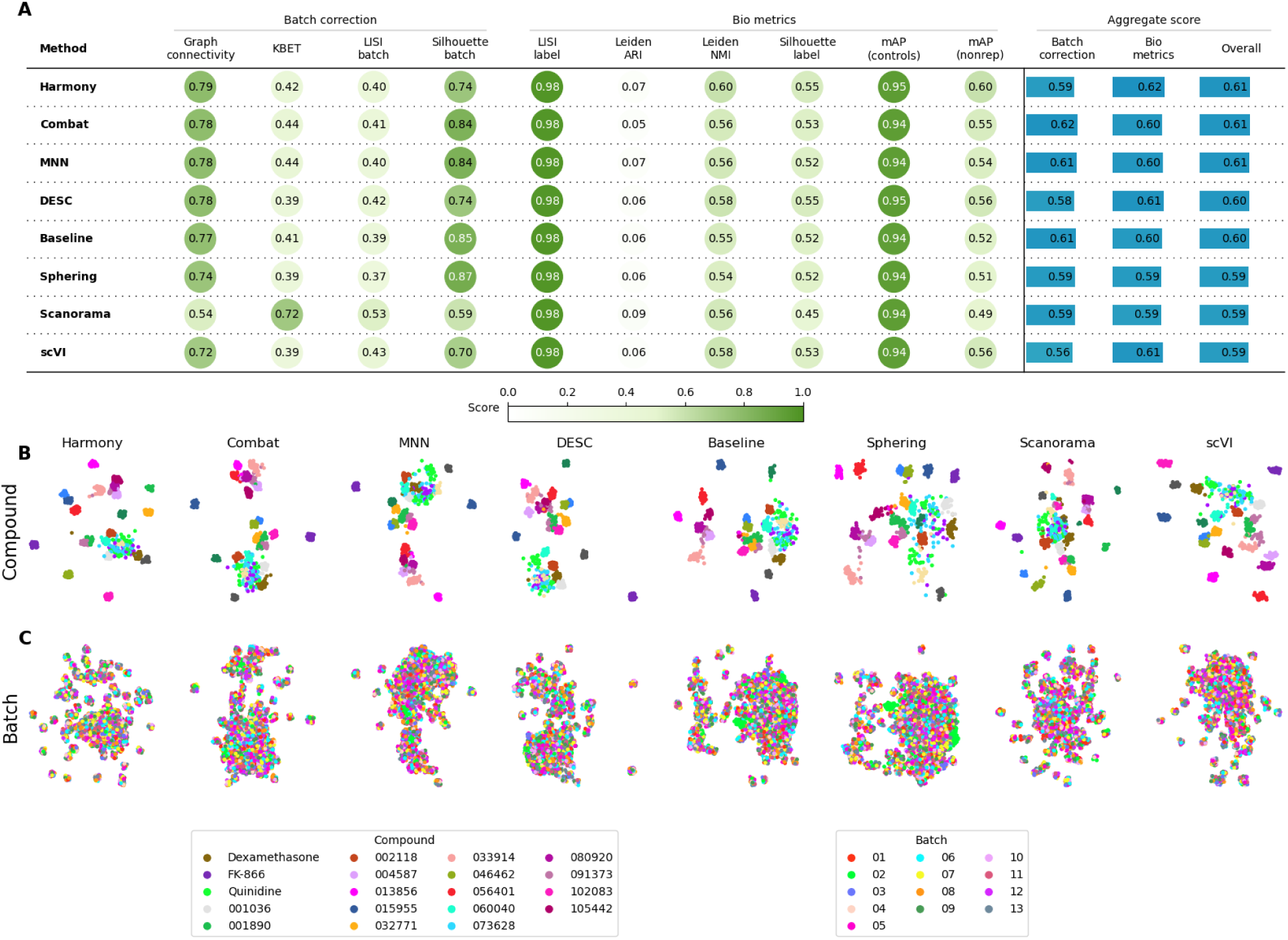
Evaluation Scenario 1. **A)** Quantitative comparison of seven batch correction methods measuring batch effect removal (four batch correction metrics) and conservation of biological variance (six bio-metrics). Metrics are mean aggregated by category. Overall score is the weighted sum of aggregated batch correction and bio-metrics with 0.4 and 0.6 weights respectively. Visualization of integrated data colored by **B)** Compound, and **C)** Batch. Left-to-right layout reflects the methods’ descending order of performance. We selected 18 out of 306 compounds with replicates in different well positions to account for position effects that may cause profiles to look similar. Alphanumeric IDs denote positive controls.

### Scenario 2

**Supplementary Figure C.**
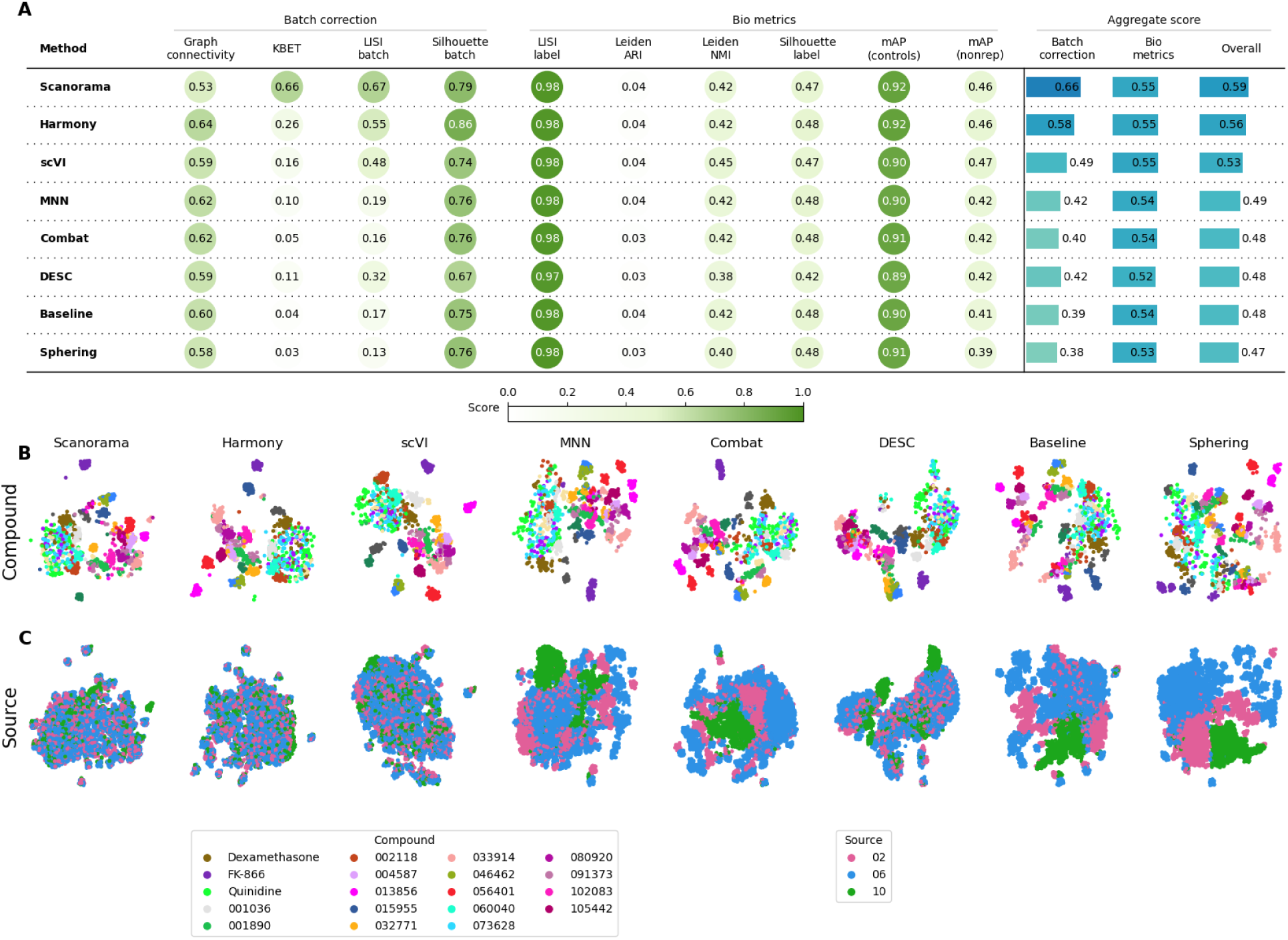
Evaluation Scenario 2. **A)** Quantitative comparison of seven batch correction methods measuring batch effect removal (four batch correction metrics) and conservation of biological variance (six bio-metrics). Metrics are mean aggregated by category. Overall score is the weighted sum of aggregated batch correction and bio-metrics with 0.4 and 0.6 weights respectively. Visualization of integrated data colored by **B)** Compound, and **C)** Source. Left-to-right layout reflects the methods’ descending order of performance. We selected 18 out of 306 compounds with replicates in different well positions to account for position effects that may cause profiles to look similar. Alphanumeric IDs denote positive controls.

### Scenario 3

**Supplementary Figure D.**
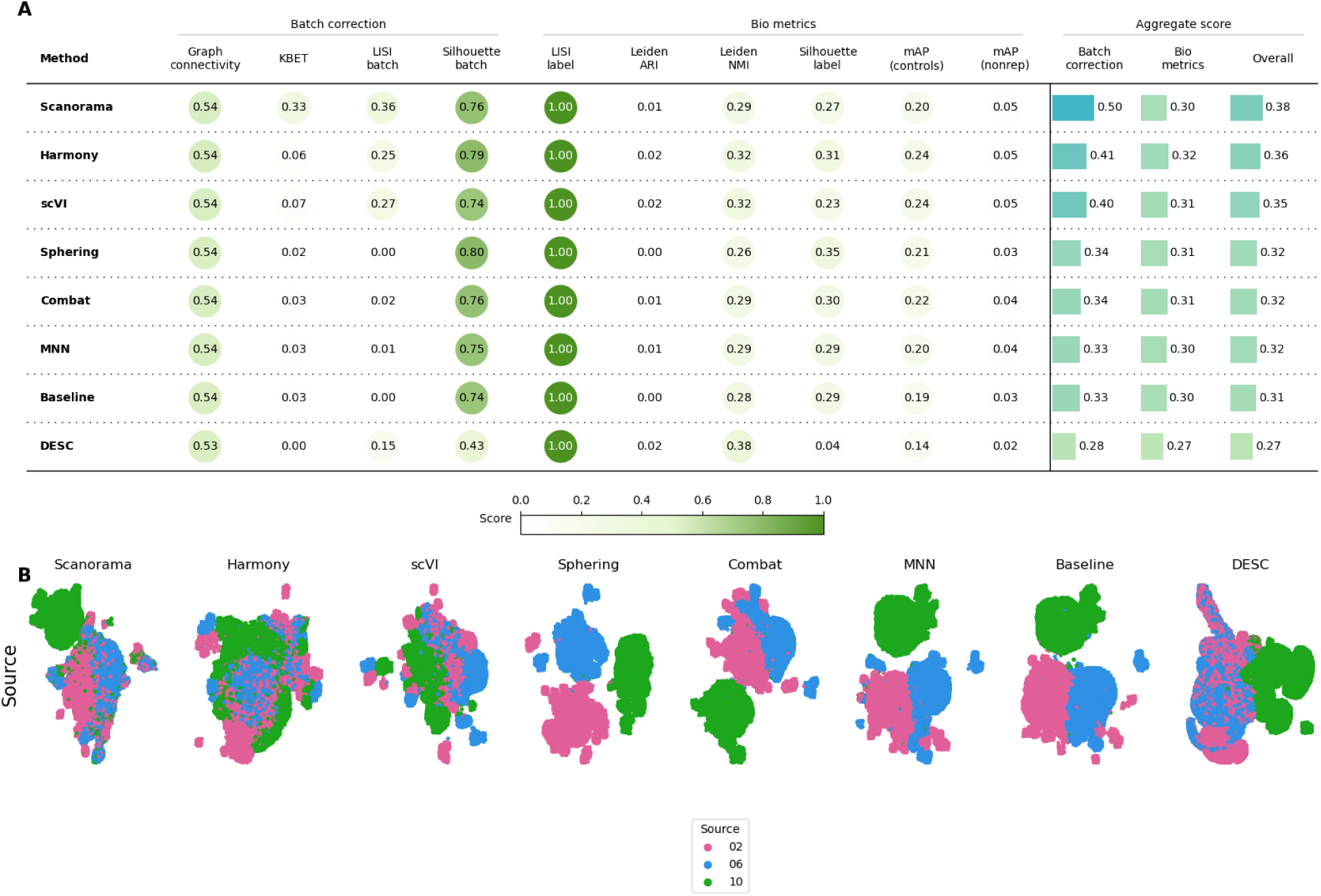
Evaluation Scenario 3. **A)** Quantitative comparison of seven batch correction methods measuring batch effect removal (four batch correction metrics) and conservation of biological variance (six bio-metrics). Metrics are mean aggregated by category. Overall score is the weighted sum of aggregated batch correction and bio-metrics with 0.4 and 0.6 weights respectively. **B)** Visualization of integrated data colored by Source. Left-to-right layout reflects the methods’ descending order of performance.

### Scenario 5

**Supplementary Figure E.**
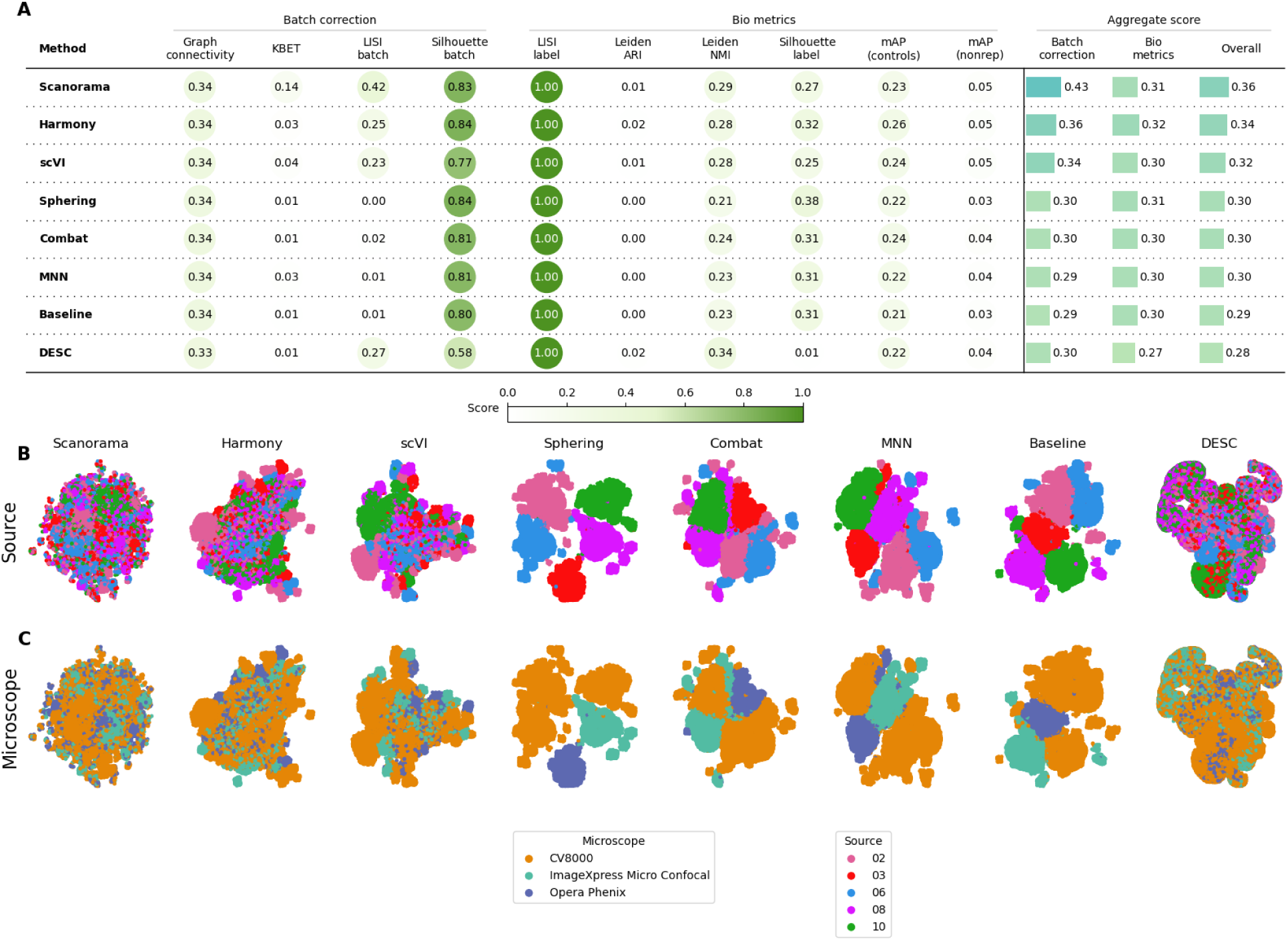
Evaluation Scenario 5. **A)** Quantitative comparison of seven batch correction methods measuring batch effect removal (four batch correction metrics) and conservation of biological variance (six bio-metrics). Metrics are mean aggregated by category. Overall score is the weighted sum of aggregated batch correction and bio-metrics with 0.4 and 0.6 weights respectively. Visualization of integrated data colored by **B)** Source, and **C)** Microscope. Left-to-right layout reflects the methods’ descending order of performance.

### Isolated compounds performance

Around 30% of the compounds of Scenario 3 are present in all three sources (sources 2, 6, and 10). We used this scenario to assess the replicate retrieval performance of sub-populations of compounds that are not shared between different batches (i.e. sources, in this setup). We used the corrected profiles from the best-performing correction method in the scenario – Harmony – to evaluate. We picked the 10,136 compounds that present in sources 2 and 6 but not in source 10 (i.e., they are isolated to sources 2 and 6). We compared the performance of this subpopulation (named as **two sources** in Sup Figure F) with the performance of a subpopulation of 23,782 compounds present in all of the three sources (named as **three sources** in Sup Figure F). Then we compute the mAP (control) score for each subpopulation, noting that we pick only the replicates from source 2 and source 6 and ignoring the replicate from source 10. We observed that the compounds that exclusively belong to **two sources** performed better than compounds present in all **three sources**, which contradicts the over-correction hypothesis. A likely explanation is that the correction task gets more difficult as there are more sources to align.

**Supplementary Figure F:**
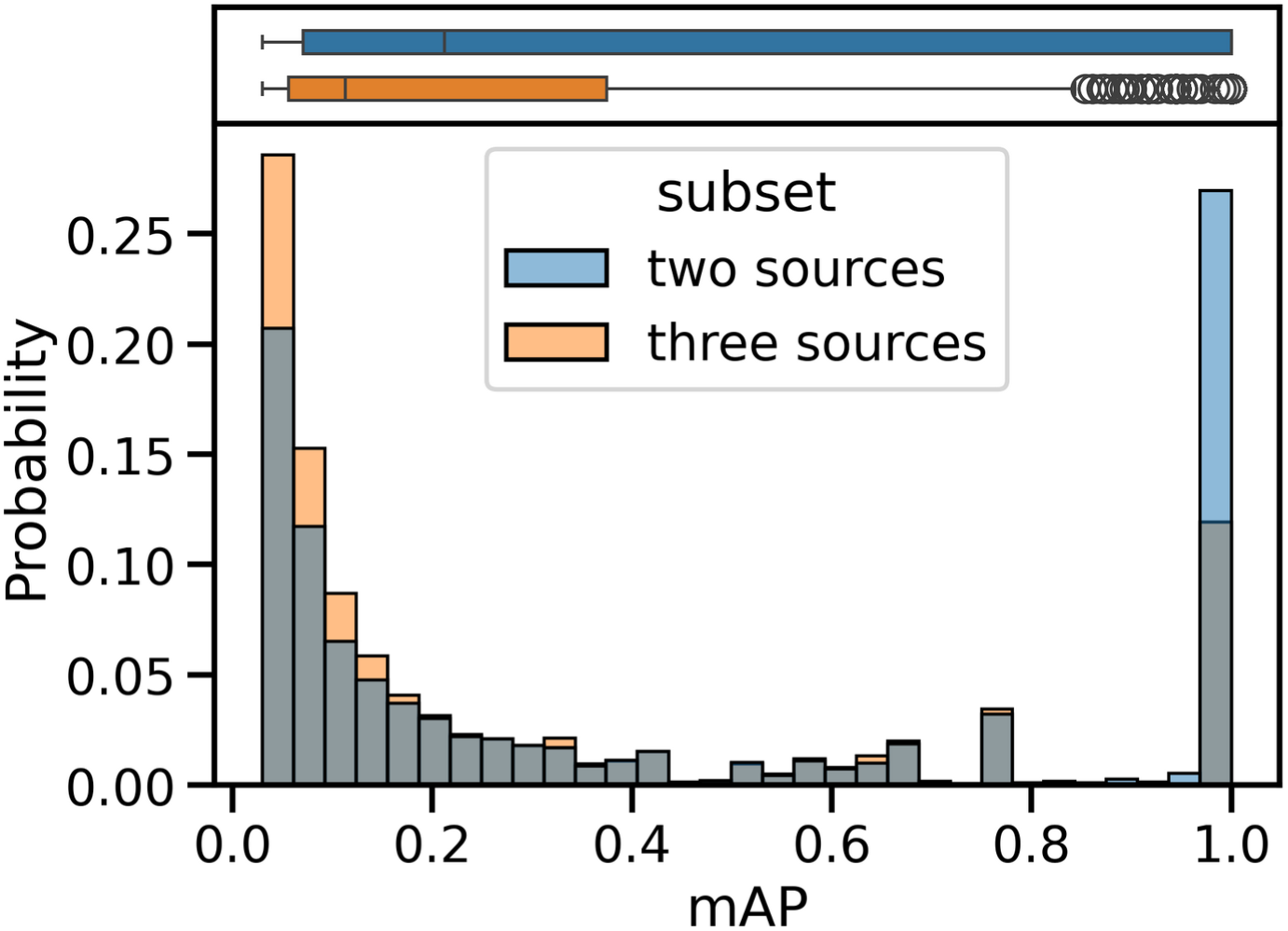
Comparison of the replicate retrieval performance (mAP) of sub-populations of compounds that are not shared between different batches. The sub-population present in only *two sources* performed better than the sub-population in *three* sources. Data extracted from the Scenario 3.

### Runtime analysis

We measured the runtime for non-gpu methods and metrics across the five scenarios on a c6i.16xlarge AWS EC2 instance equipped with 64 cores and 128GB of RAM. A log-log plot of the results (Sup Figure G) reveals a linear-like trend, suggesting a power-law relationship between runtime and sample size. Extrapolating this trend, applying Harmony (a top-performant method) at the single-cell level (Sup Table A) would be prohibitively time-consuming: approximately 2.6 hours for a single plate, 33 hours for a single batch, and 11 days for a single source (there are 13 sources in the full JUMP Cell Painting dataset). Furthermore, loading such a source would require 2.7 TB of memory.

**Supplementary Figure G:**
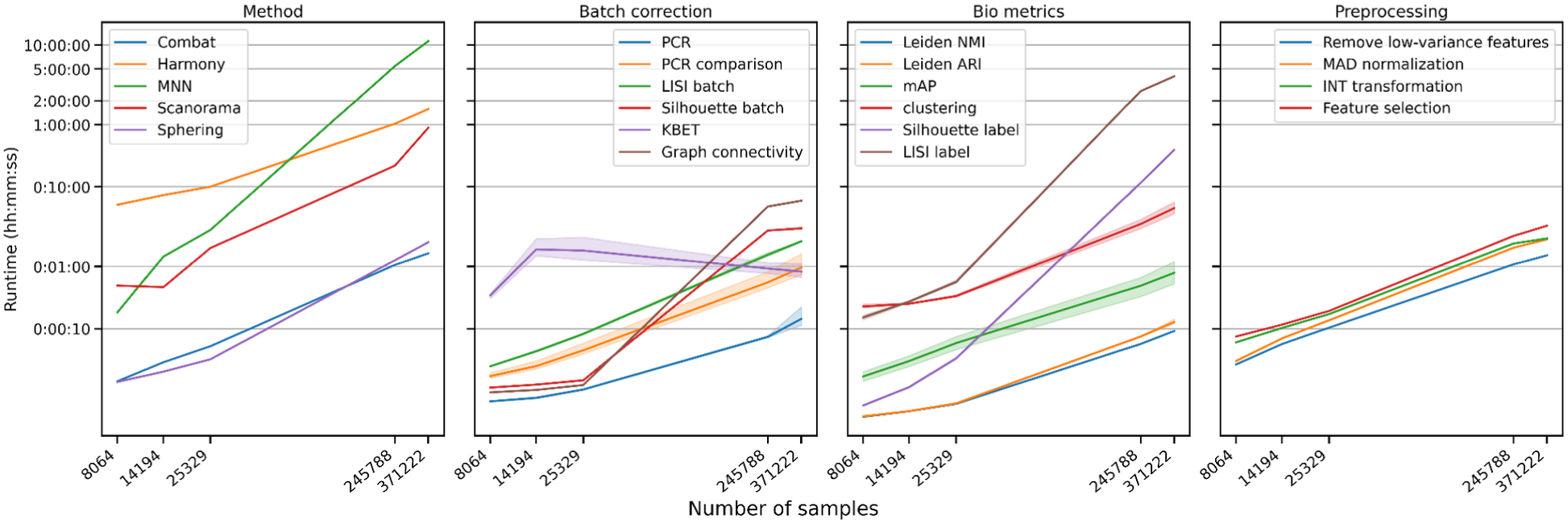
Runtime analysis of methods, metrics, and preprocessing steps for all the scenarios. Scenario 1, 2, and 4 (first three ticks in the x-axis) have *JUMP-Target-2-Compound* plates only with ∼300 unique compounds. Scenarios 3 and 5 (last two ticks in the x-axis) have Production plates with ∼80,000 unique compounds. Both axes are log-scaled. KBET runtime trend is constant because it only evaluates compounds with more than 15 replicates. The Clustering step is required for LISI, NMI, ARI, Graph connectivity, and KBET.

**Supplementary Table A:**
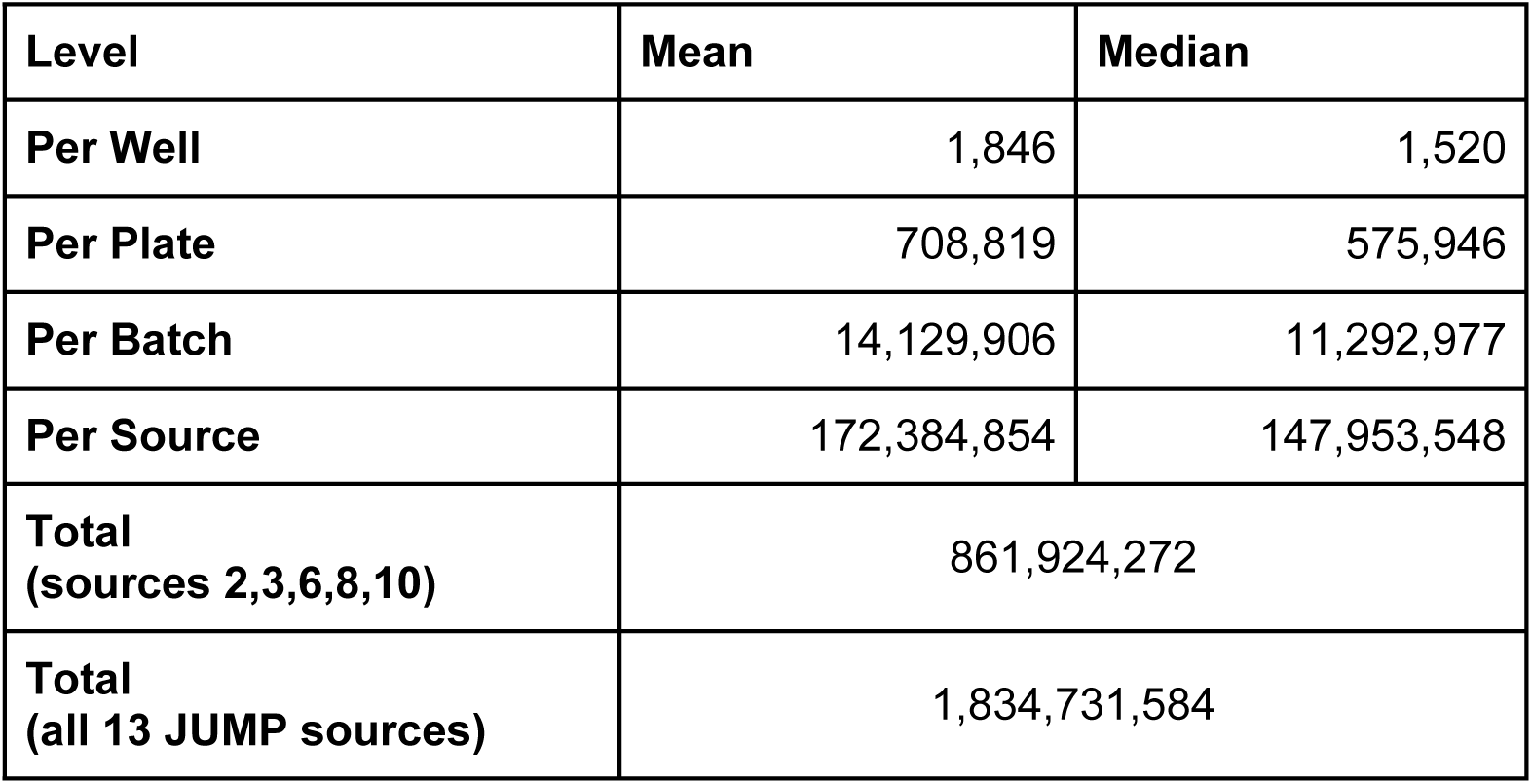
Count of single cells at different levels in the JUMP CP Dataset.

### Performance distribution

**Supplementary Figure H:**
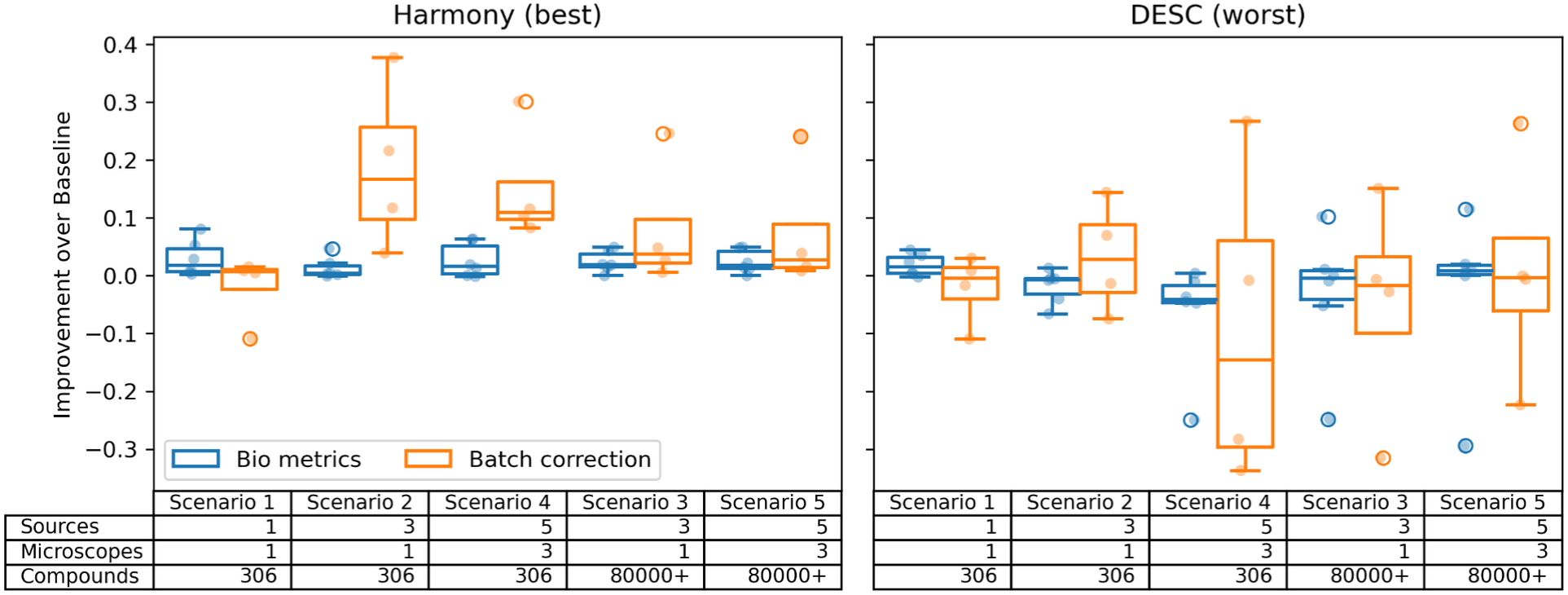
Comparison of best and worst batch correction methods, reflecting the variability of the performance with respect to the complexity of the scenarios (scenarios are sorted by overall mean score).

**Supplementary Figure I:**
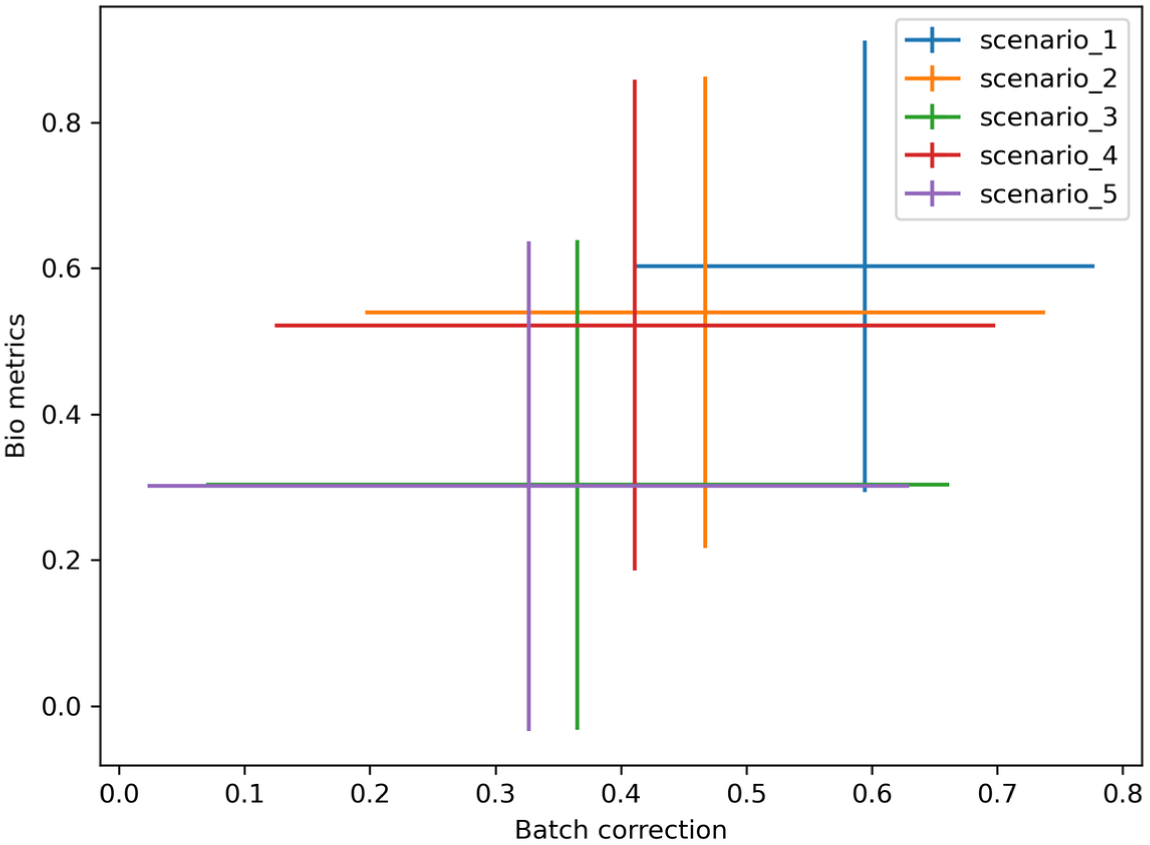
Scatter plot of the mean Batch correction and bio-metrics for all the tested methods across the five scenarios, reflecting the increasing difficulty of scenarios. Bars represent one standard deviation in the respective axis.

### Implementation notes

- For scVI, we shifted the data to the feasible space. (i.e. transform each feature 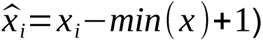
- The nature of the image-based profile data involving low number of replicates and high number of compounds limits kBET, which relies on a higher (>15) number of samples per biological concept.
- mAP is the only metric able to capture the performance of the models when there are as few as only two replicates of a compound.
- We optimize the preprocessing pipeline based on a mAP.
- We adjust DESC convergence hyperparameters to avoid collapsed representations (default parameters converged to vectors with only −1, 1 values (output from a tanh activation) )
- We increase the number of Harmony clusters from 50 to 300 and iterations from 10 to 20.
- We increase the number of latent dimensions in scVI from 10 to 30.
- Combat implementation from scanpy has no hyperparameters.
- We use the default hyperparameters for MNN (neighbor size=20), Scanorama (KNN=20, alpha=0.1, sigma=15) and scVI (num_units=128, dropout=0.1).

## Footnotes

1 https://github.com/theislab/scib

